# Chromatin state dynamics during the *Plasmodium falciparum* intraerythrocytic development cycle

**DOI:** 10.1101/2025.08.22.671872

**Authors:** Alan S. Brown, Manuel Llinás, Shaun Mahony

## Abstract

The interdependence of chromatin states and transcription factor (TF) binding in eukaryotic genomes is critical for the proper regulation of gene expression. In this study, we explore the connection between TFs and chromatin states in the human malaria parasite, *Plasmodium falciparum*, throughout its 48-hour asexual intraerythrocytic developmental cycle (IDC). Most *P. falciparum* genes are expressed in a periodic manner during the IDC, accompanied by dynamic shifts in histone modifications and chromatin accessibility. Leveraging genome-wide profiles of chromatin accessibility, histone modifications, and Heterochromatin Protein 1 (HP1) occupancy, we characterize chromatin state dynamics during the IDC. Our results indicate that several chromatin states remain stable throughout the lifecycle, while others are dynamic and are linked to gene activation or repression. We further characterize chromatin state dynamics at the genome-wide DNA binding sites for a selection of *Plasmodium* TFs, allowing us to group TFs according to their chromatin preferences. By correlating changes in chromatin accessibility, histone modifications, and TF binding, we provide a global overview of the chromatin state dynamics that coordinate *P. falciparum* asexual blood stage development.

## Introduction

Methods for identifying combinatorial patterns of histone modifications, or “chromatin states”, have been applied to characterize regulatory elements across several multicellular eukaryotic species, including *Arabidopsis thaliana*, *Caenorhabditis elegans*, *Drosophila melanogaster*, mouse, and human (Ernst et al., 2011; Ernst & Kellis, 2012, 2017; Kharchenko et al., 2011, 2011; Liu et al., 2011; Roudier et al., 2011; Vu & Ernst, 2023). By integrating multiple datasets into categorical variables representing co-occurring patterns of histone modifications, chromatin states distill complex epigenetic information into more easily interpretable regulatory categories. Chromatin state analysis approaches have been effectively used in large scale genomic research projects to annotate key regulatory features across many cell types and conditions (Vu & Ernst, 2022, 2023). For example, ENCODE and other mammalian epigenomic consortium studies used ChromHMM to define chromatin states by applying a hidden Markov model to binarized chromatin signals and annotate potential regulatory regions (ENCODE Consortium, 2004, 2012; Ernst & Kellis, 2012, 2017; Kundaje et al., 2015; Vu & Ernst, 2022, 2023). Similarly, Segway, a dynamic Bayesian network-based approach for segmenting genomic data, was also deployed by ENCODE (ENCODE Consortium, 2004, 2012; Hoffman et al., 2012, 2013). In other systems such as human and mouse hematopoiesis, the integrative and discriminative epigenome annotation system (IDEAS), which uses a 2D segmentation approach to define consistent chromatin states across genomic locations and cell types, has been used to integrate epigenomics datasets (Hardison et al., 2020; Xiang et al., 2021; Zhang et al., 2016; Zhang & Hardison, 2017; Zhang & Mahony, 2019).

Chromatin state analyses have not been widely used to characterize the regulatory features in single-celled eukaryotes such as the human malaria parasite, *Plasmodium falciparum*. The lifecycle of *P. falciparum* includes a 48-hour asexual intraerythrocytic developmental cycle (IDC), during which the parasite transitions through three distinct morphological stages – rings, trophozoites, and schizonts – inside a red blood cell before egressing as merozoites to invade new red blood cells (Bannister & Mitchell, 2003). Gene expression is highly dynamic during the IDC, with most genes displaying a periodic expression pattern that correlates with specific stages of development (Bozdech et al., 2003; Chappell et al., 2020; Le Roch et al., 2003; van Biljon et al., 2019). While several studies have characterized individual histone modifications, histone variants, chromatin accessibility, transcription factor binding, and transcript abundance profiling during the IDC (Bartfai et al., 2010; Bunnik et al., 2018; Carrington et al., 2021; Gupta et al., 2013; Jiang et al., 2013; Karmodiya et al., 2015; Ruiz et al., 2018; Salcedo-Amaya et al., 2009; Shang et al., 2022; Tang et al., 2020; Toenhake et al., 2018), our understanding of how combinations of these chromatin features relate to regulatory dynamics remains incomplete. A single previous study has profiled chromatin states based on three histone modifications in parasites that were transitioning through the merozoite stage (Reers et al., 2023), but this did not fully describe chromatin state dynamics throughout the IDC. We also previously demonstrated that *P. falciparum* TF binding site selection is impacted by chromatin features (Bonnell et al., 2024). Thus, a more complete characterization of chromatin state dynamics is crucial for understanding the tightly regulated asexual developmental cycle of *Plasmodium* parasites.

The *P. falciparum* genome has several unique features that make it uncertain whether chromatin state principles derived from other species are directly transferable. Its genome is extremely A/T-rich, with an average A/T content of 80% and up to 90% in non-coding regions (Gardner et al., 2002; Otto et al., 2018; Silberhorn et al., 2016). Although *Plasmodium* parasites encode all of the canonical histones (Miao et al., 2006; Trelle et al., 2009) except the H1 linker (Gardner et al., 2002; Sullivan et al., 2006), the *Plasmodium* histone H2A sequences are somewhat divergent from other eukaryotes (Watzlowik et al., 2021). Other properties of *Plasmodium* nucleosomes also differ from those of humans and yeast, including reduced stability under varying conditions, and decreased effectiveness at binding GC-rich DNA (Silberhorn et al., 2016). Importantly, some histone modifications have atypical associations with regulatory activities in *P. falciparum* compared to other eukaryotes. For instance, H3K36me3, which typically marks active transcription in *Drosophila*, mouse, and humans (Bell et al., 2007; Schwartz et al., 2009), is instead associated with gene silencing in *P. falciparum* (Connacher et al., 2021; Jiang et al., 2013). H3K4me1 has been associated with both poised chromatin states and active regulatory elements in *Plasmodium* (Karmodiya et al., 2015; Ubhe et al., 2017), whereas in animal genomes it primarily marks distal enhancer elements (Creyghton et al., 2010; Heintzman et al., 2009). These differences in chromatin associations highlight the need for a *Plasmodium*-specific understanding of chromatin state dynamics.

Studies of individual epigenomic features point to a highly dynamic chromatin landscape across different stages of the *P. falciparum* IDC. For example, chromatin accessibility increases significantly after the ring stage and peaks during the trophozoite or early schizont stages, correlating with enhanced gene expression (Ponts et al., 2010; Ruiz et al., 2018; Toenhake et al., 2018). H3K4 methylation increases after the ring stage, while H3K27 acetylation is highest during the ring stage, and H3K9 acetylation predominates in the trophozoite stage (Bartfai et al., 2010; Coetzee et al., 2017; Gupta et al., 2013). The highly dynamic expression program is tightly controlled by a relatively small number of transcription factors (TFs); there are only 73 predicted TFs encoded in the *Plasmodium* genome (Balaji et al., 2005; Bischoff & Vaquero, 2010; Coulson et al., 2004), including 30 members of the apicomplexan APETALA2 (ApiAP2) family that have no known homologs in humans (Singhal et al., 2024). Understanding how this limited TF repertoire interacts with the dynamic chromatin landscape is essential for deciphering the regulatory mechanisms that drive developmental transitions in *Plasmodium* parasites.

In this study, we comprehensively characterize chromatin state dynamics throughout the *P. falciparum* IDC by integrating data on chromatin accessibility, histone modifications, and Heterochromatin Protein 1 occupancy. We further analyze these chromatin states at the binding sites of key ApiAP2 transcription factors, allowing us to group TFs according to chromatin state preferences. By correlating changes in chromatin accessibility, histone modifications, and TF binding, we provide insights into the gene regulatory network that coordinates asexual blood stage development of the malaria parasite during infection of the human host.

## Results

### Defining chromatin states during the *P. falciparum* intraerythrocytic development cycle

To define chromatin regulatory landscapes during the *P. falciparum* IDC, we compiled epigenomic data from multiple published sources (see **Methods**). Since these experiments were generated by individual labs rather than a coordinated effort, only a subset of signals has been profiled across consistent developmental timepoints. We therefore focused on the three main stages of the IDC: ring stage (10 hours post invasion (hpi)); trophozoite stage (30 hpi); and schizont stage (40 hpi). Our analysis included seven histone modifications/variants with data available for all three stages: H3K4me1, H3K4me3, H3K9me3, H3K9ac, H3K18ac, H3K27ac, and H2A.Z, along with ATAC-seq data measuring chromatin accessibility. We also incorporated ChIP-seq data for Heterochromatin Protein 1 (HP1), a known indicator of heterochromatic regions.

We used ChromHMM (Ernst & Kellis, 2012) to identify chromatin states across the *P. falciparum* genome at a resolution of 200bp bins, determining 11 states to be optimal after testing various state numbers (see **Methods**). Our results identified the probabilities of observing each epigenomic signal in regions covered by each of the 11 states (**Fig. 1A**). We also determined the relative chromatin state enrichment at various genomic annotations (**Fig. 1B**), their genome-wide prevalence (**Fig. 1C**), and their occurrence at genes (**Fig. 1D**) across all three developmental timepoints.

**Figure 1:**
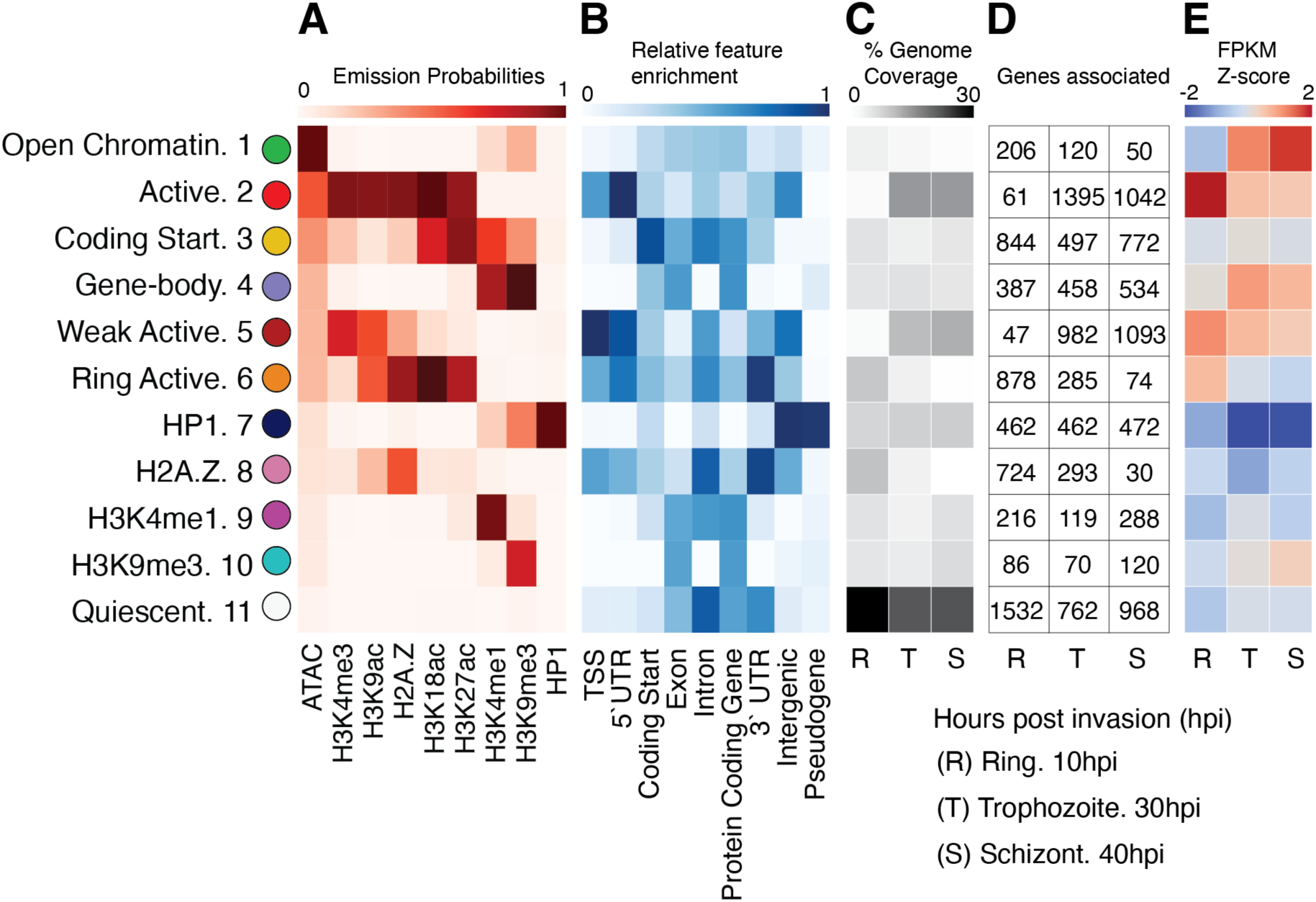
Chromatin states in the *P. falciparum* IDC and associated features. **A)** Hidden Markov Model emission probabilities for each of the 11 chromatin states discovered by ChromHMM. **B)** Relative enrichment of each state at a selection of genomic features. **C)** The genomic coverage of each state at each timepoint: R= ring stage (10hpi), T= trophozoites (30hpi), and S= schizont stage (40hpi). **D)** The number of genes for which each state is located at the start of the coding sequence at each timepoint. **E)** Z-score values for the average gene expression value (RNA-seq FPKMs) per chromatin state.

Several identified chromatin states parallel those observed in other eukaryotic systems, including those characterized in the human genome by the ENCODE and ROADMAP consortia (ENCODE Consortium, 2004, 2012; Kundaje et al., 2015). Chromatin state 2, which is characterized by a combination of chromatin accessibility, H3K4me3, H2A.Z, and various histone acetylation marks, is highly enriched at transcription start sites (TSSs), similar to active promoter states in vertebrates and *Drosophila* (Brown & Bachtrog, 2014; Ernst & Kellis, 2015, 2017; Vu & Ernst, 2023). Other states that include H3K4me3 and/or H2A.Z marks (chromatin states 5, 6, and 8) also show associations with TSSs and 5’ untranslated regions (5’ UTRs). Chromatin state 3, which is enriched for H3K4me1, H3K27ac, and ATAC-seq signals, resembles enhancer-associated signatures in vertebrates (Ernst & Kellis, 2015, 2017). However, unlike vertebrate genomes, where these marks identify distal enhancers, the *P. falciparum* chromatin state 3 is more highly associated with coding start regions and gene bodies than with intergenic regions. Chromatin state 3 may thus be analogous to a H3K4me1-enriched chromatin state that is observed flanking TSSs in vertebrate genomes (Ernst & Kellis, 2015, 2017). Chromatin state 11, which is largely depleted of any tested epigenomic signals, resembles the “quiescent” chromatin state that covers up to 80% of the human genome (Ernst & Kellis, 2017; van der Velde et al., 2021). This state is also the most frequently occurring in *P. falciparum*, appearing in 25-30% of the measured genomic windows (**Fig. 1C**). Chromatin state 1, displaying only ATAC-seq enrichment, is seen relatively rarely in *P. falciparum*, although it is analogous to a similar chromatin state observed in the human genome (Vu & Ernst, 2022).

While several *P. falciparum* chromatin states have analogs in vertebrate systems, the H3K9me3-associated states show some unique properties. In the human genome, H3K9me3 typically marks repressed heterochromatin. For example, chromatin state 7 in our analysis is characterized by enrichment of both H3K9me3 and HP1 and likely represents heterochromatic regions. However, chromatin state 4 displays enrichment of both H3K9me3 and H3K4me1, without HP1, and is associated with exonic sequences. Similarly, states displaying only H3K4me1 (chromatin state 9) or only H3K9me3 (chromatin state 10) are also associated with exons. Thus, *P. falciparum* may use H3K9me3 and H3K4me1 to mark genomic processes in a manner that differs from vertebrate genomes.

Examining the relationships between chromatin states and gene expression, we find that three of the four TSS-associated states (states 2, 5, and 6) correlate with higher expression levels (**Fig. 1E**), consistent with “active” TSS states in human. The exception is the H2A.Z-enriched chromatin state 8, which is associated with lower expression levels and is differentiated from other TSS-associated states by a relative lack of histone acetylation signals. Interestingly, the prevalence of TSS-associated states varies dramatically across the IDC. Chromatin states 6 and 8 are common during the ring stage (11.6% and 12.1% of genomic regions, respectively) but become rare at later timepoints. Conversely, chromatin states 2 and 5 are rare during the ring stage, but each mark the start of over 1,000 genes during trophozoite and schizont stages.

As expected, the putative heterochromatin state (state 7) correlates with repressed expression (**Fig. 1E**) and is stably associated with approximately 10% of the genome throughout the IDC. This state is primarily found in telomeric, subtelomeric, and centromeric regions, and encompasses most of the clonally variant *var* and *rifin* genes involved in immune system evasion through antigenic variation (Pickford et al., 2021). Most of these genes are constitutively repressed, with only a subset (variable across a population of cells) gaining high expression levels (Tripathi et al., 2022). Finally, chromatin state 1 (ATAC-seq only) is associated with low levels of expression in the ring stage, but higher levels in trophozoite and schizont stages, potentially indicating a “poised” regulatory signature, though its prevalence decreases during the life cycle (from 4.8% genome coverage in ring stage to only 1.8% in schizonts).

### Chromatin states are temporally dynamic during the *P. falciparum* IDC

Previous work in human and model organisms has demonstrated that chromatin states can change rapidly over developmental time, reflecting underlying dynamics in gene expression and regulatory mechanisms (Chang et al., 2023; Mattout et al., 2015; Yadav et al., 2018). Building on our stage-specific chromatin state analyses, we evaluated the degree to which chromatin states change at individual genomic bins (200bp) throughout the *P. falciparum* IDC (**Fig. 2A**). Approximately two-thirds of all genomic bins display at least one change in chromatin state across the three developmental stages examined (**Fig. 2B**), indicating highly dynamic chromatin behavior during the *P. falciparum* IDC.

**Figure 2:**
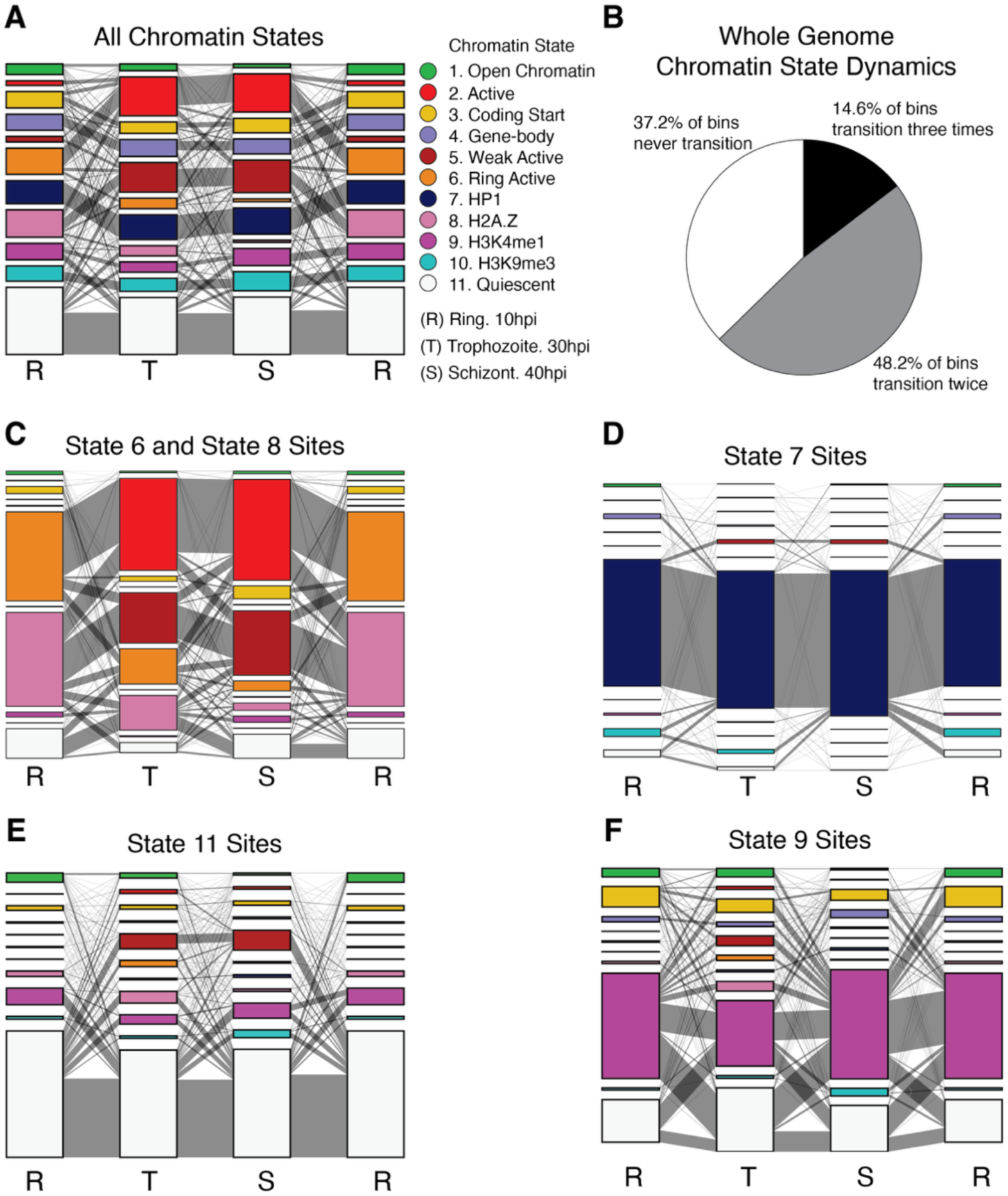
Chromatin state dynamics during the *P. falciparum* IDC. **A)** Chromatin state transitions across all 200bp genomic bins. Stacked bar graphs show the relative proportions of the 11 chromatin states during ring (R), trophozoite (T), and schizont (S) stages. The width of connecting gray lines between stages represents the number of bins transitioning between each pair of chromatin states. Ring stage is displayed twice to illustrate the full developmental cycle from schizont back to ring stage. The visualization format is maintained in panels C)-F), with each panel focusing on a specific subset of genomic bins to illustrate different aspects of chromatin state dynamics. **B)** Proportion of genomic bins that change states across the IDC. “Never transition” indicates bins that maintained the same state across all 3 timepoints. “Transition twice” represents bins that change to a different state and then revert to their original state. “Transition three times” indicates bins that display a different state at each of the three timepoints. **C)** Chromatin state transitions for bins displaying the “ring active” (state 6) or “H2A.Z” (state 8) chromatin states at any timepoint. **D)** Chromatin state transitions for bins displaying the heterochromatin-associated state 7. **E)** Chromatin state transitions for bins that are labeled quiescent state in at least one timepoint. **F)** Chromatin state transitions for bins displaying the H3K4me1-associated state 9.

As previously noted, the various TSS-associated chromatin states (2, 5, 6, and 8) each undergo shifts in global abundance during the IDC. Investigating transitions to or from states 6 or 8, we observed a clear shift in chromatin profiles between the ring stage and the two later trophozoite and schizont stages (**Fig. 2C**). Specifically, states 6 and 8 have a clear progression into states 2 and 5 at later timepoints. Of the genomic bins displaying chromatin state 6 during the ring stage, 71.9% transition to state 2 in the trophozoite stage. Similarly, bins with the H2A.Z-enriched state 8 at ring stage transition to state 5 (41.9%) or state 2 (29.2%) in trophozoites. As previously mentioned, state 8 is not associated with actively transcribed genes, possibly reflecting the overall lower transcriptional activity in the ring stage (Bozdech et al., 2003; Llinás et al., 2006).

The rapid decrease in global prevalence of states 6 and 8 and their transitions can be explained by a global increase in the histone mark H3K4me3. The main differences between states 2 and 6 and between states 5 and 8 are the relative levels of H3K4me3 in their respective emission probabilities (**Fig. 1A**). Our analyses use ChIP-seq data showing globally low levels of H3K4me3 in ring stage parasites but higher levels in trophozoites and schizonts (Bartfai et al., 2010). Other studies corroborate the low level of H3K4me3 in ring stage parasites (Coetzee et al., 2017; Gupta et al., 2013; Reers et al., 2023). Thus, part of the dynamic behavior we observe in chromatin states may reflect a shift in the types of histone modifications present on the *P. falciparum* genome during the IDC.

The persistence of chromatin states over time varies considerably between states. The most stable state is the heterochromatin-associated state 7 (**Fig. 2D**). Of the genomic bins associated with chromatin state 7 at any timepoint, 79% maintain state 7 across all three timepoints. These stable state 7 loci exist primarily in the sub-telomeric and centromeric regions, which remain consistently heterochromatic. Most genes in these regions are repressed, although a small number of clonally variant genes are expressed at high levels non-uniformly across the parasite population (Tripathi et al., 2022). The second-most stable state is the quiescent state 11, with 38.8% of quiescent sites remaining quiescent at all three timepoints. Examining each transition between timepoints reveals that an average of 70.5% of the sites annotated with state 11 remain state 11 in the next time point (62.4% from 10hpi to 30hpi, 72.5% from 30hpi to 40hpi, and 76.6% from 40hpi back to 10hpi) (**Fig. 2E**). Of the 29.5% that transition from the quiescent state to some other state, the most common target state is state 9 when transitioning from trophozoite to schizont and schizont to ring. During the ring stage, the most common transition target from the quiescent state is state 5, followed by state 8, and then state 9. This pattern generally aligns with the broader trend of ring-stage chromatin states shifting to state 5. However, it diverges in the proportion that changes to H2A.Z-enriched state 8, which decreases overall from ring to trophozoite stages.

In contrast to the stable heterochromatin state, and aside from the TSS-associated states that display large global prevalence changes, the most dynamic state is the H3K4me1-associated state 9. Of the over 13,000 bins that are state 9 in at least one time point, only about 10% remain in state 9 at all measured timepoints. Among bins that change across time, an average of 32.5% of state 9 bins transition to the quiescent state in the subsequent point (43.6% from 10hpi to 30hpi, 32.2% from 30hpi to 40hpi, and 21.8% from 40hpi back to 10hpi) (**Fig. 2F**). Overall, our findings demonstrate that temporal changes in chromatin states vary dramatically from highly stable and therefore unlikely to result in transcriptional changes, to large changes that can modulate gene expression during the IDC.

### Chromatin state dynamics are associated with gene expression dynamics

While we find that specific chromatin states correlate with either elevated or reduced gene expression levels at individual timepoints (**Fig. 1E**), both chromatin states and transcriptional activity exhibit highly dynamic and periodic patterns during parasite development. We therefore investigated how temporal changes in chromatin states relate to general shifts in gene expression patterns across the parasite’s lifecycle. To analyze this relationship, we assigned a single representative chromatin state to each gene. Based on evidence that histone modifications near the start codon are most predictive of expression levels (Read et al., 2019), we assigned each gene the most prevalent chromatin state within a 600bp window surrounding its start codon (see **Methods**). We next measured expression distributions for genes that change chromatin states between consecutive timepoints and categorized them according to the chromatin state gained (**Fig. 3**). Interestingly, a subset of genes do not change chromatin states between consecutive pairs of timepoints (**Supplementary Fig. 1**) and others display no change in chromatin states throughout the IDC (**Supplementary Fig. 2**).

**Figure 3:**
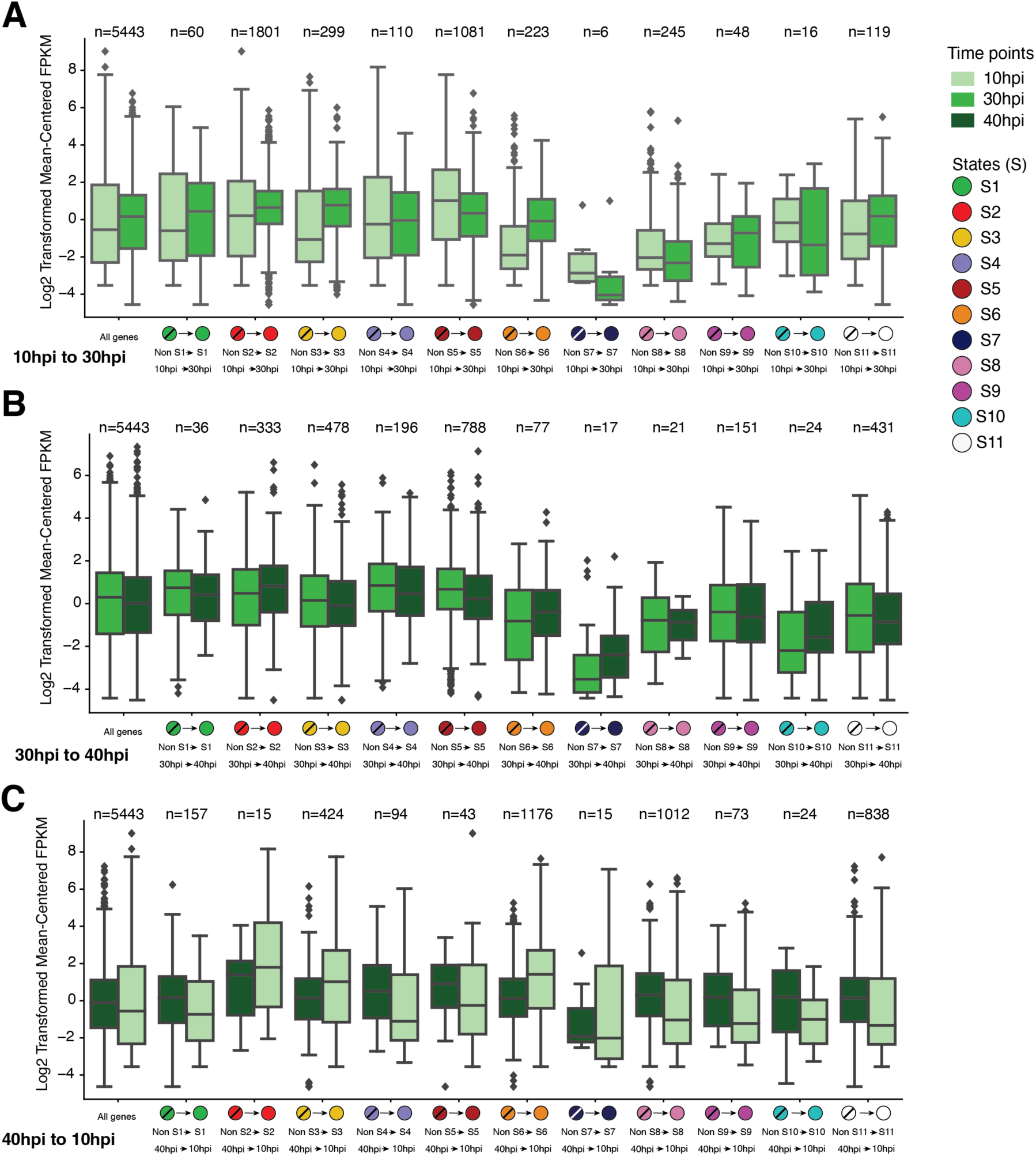
Associations between chromatin state transitions and expression dynamics at genes that change states between consecutive pairs of timepoints. Expression boxplots display the distributions of log_2_-transformed mean-centered FPKM normalized RNA-seq values for groups of genes that display indicated transitions from other states into the labeled state at the labeled timepoints. **A)** Chromatin state transitions and expression values from 10hpi to 30hpi. **B)** Chromatin state transitions and expression values from 30hpi to 40hpi. **C)** Chromatin state transitions and expression values from 40hpi back to 10hpi. See **Supplementary Figure 1** for a similar plot showing groups of genes that display no change in chromatin states between timepoints.

Genes transitioning into the acetylation-rich active states 2 or 6 display increases in average expression across all three timepoint transitions (**Fig. 3**). This expression gain is particularly striking for the large group of 1,176 genes transitioning into state 6 at the ring stage (i.e., from 40hpi to 10hpi). Transitions into state 3, another acetylation rich state, also correlated with gains in gene expression. Surprisingly, while “weak active” state 5 is generally associated with higher expression levels (**Fig. 1E**), transitions into this state were associated with reductions in expression across all three timepoint transitions (**Fig. 3**). This suggests that gaining chromatin state 5 might represent a reduction in expression levels at otherwise highly expressed genes.

Transitions to the H3K9me3-associated states 4, 7, or 10 generally correlated with reductions in expression (**Fig. 3**). However, this trend was muted, or even reversed, during the transition into schizont stage (i.e., 30hpi to 40hpi). It should also be noted that relatively few genes gain state 7 or 10 during any transition. Transitions into open chromatin state 1, H3K4me1-associated state 9, or quiescent state 11 display mixed, stage-specific associations with expression changes. On average, a gain of one of these three states is associated with upregulation in trophozoites (i.e., 10hpi to 30hpi), a maintenance of expression levels in schizonts (i.e., 30hpi to 40hpi), and downregulation in rings (i.e., 40hpi to 10hpi). While these associations might point to context-dependent regulatory interactions between chromatin signals and expression, they may also reflect background associations as average expression levels across all genes display similar stage-specific dynamics.

To illustrate associations between chromatin states and gene expression dynamics for individual genes, we focused on a subset of genes that were selected based on their expression dynamics at different stages of the IDC (Kafsack et al., 2012) (**Fig. 4**). From this set, we highlight three representative examples (**Fig. 4B**) that peak in expression at different stages of the IDC.

**Figure 4:**
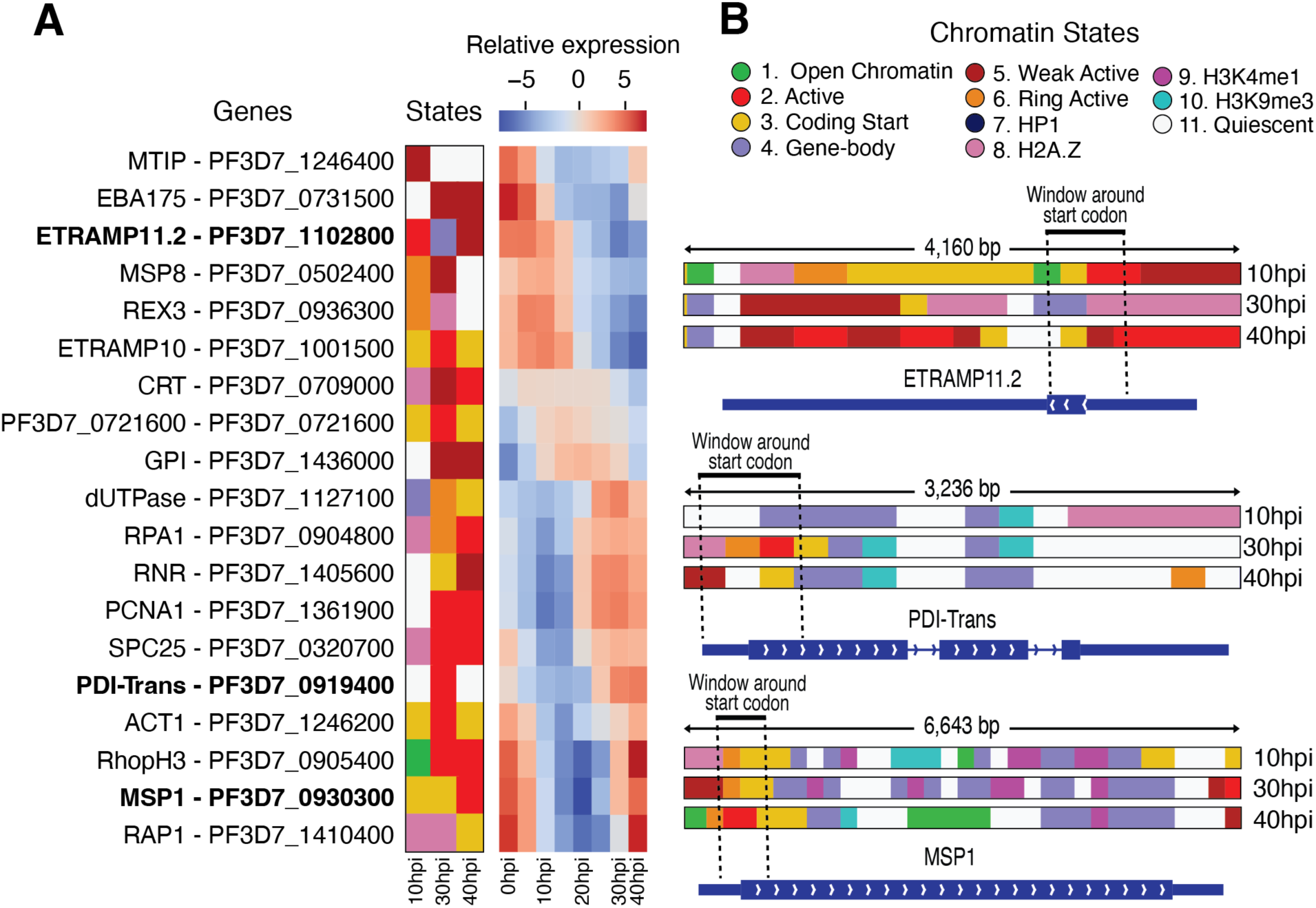
Associations between chromatin state transitions and expression dynamics for a selection of *P. falciparum* genes. **A)** Representative chromatin states at start codons at three timepoints (left) and expression heatmap from timepoints throughout the IDC (right; RNA-seq data sourced from (Toenhake et al., 2018)) for each of 19 selected genes. **B)** Chromatin states for the three timepoints at ETRAMP11.2, PDI-Trans, and MSP1 gene loci. Thick blue bars at each gene indicate exonic regions, thin blue bars indicate UTR regions and chevrons indicate the orientation of transcription.

Early-transcribed membrane protein 11.2 (ETRAMP11.2, PF3D7_1102800) is most highly expressed during the ring stage, though its expression begins in late schizonts (Spielmann et al., 2003; Toenhake et al., 2018). The chromatin profile at its start codon shows active state 2 during the ring stage, transitioning to less active state 4 in trophozoites, and then to weak active state 5 in schizonts, aligning well with its expression pattern.

The protein disulfide-isomerase PDI-Trans (PF3D7_0919400), which facilitates proper protein folding (Mahajan et al., 2006), is maximally expressed in trophozoite and schizont stages. Its chromatin states transition from quiescent state 11 in ring stage to active state 2 in trophozoites and back to quiescent state 11 in schizonts. Closer examination reveals some weak active state 5 and coding start state 3 signals near the start codon in schizonts, while the increasing presence of state 4 from trophozoites to schizonts may indicate reduced transcription in the subsequent developmental stages.

Merozoite surface protein 1 (MSP1) (PF3D7_0930300) is one of the most highly abundant proteins on the merozoite surface and as such is a promising vaccine candidate (Child et al., 2010; Das et al., 2015; James et al., 2006). MSP1 expression begins in the schizont stage while the parasite is still developing inside the host cell. Interestingly, the *msp1* gene displays the expression-neutral coding start state 3 during ring and trophozoite stages before transitioning to active state 2 in schizonts, coinciding with its rapid increase in expression. The RNA-seq data shows expression beginning to increase in trophozoites, corresponding with the appearance of weak active state 5 near the start codon. Overall, these examples demonstrate a positive correlation between expression dynamics for genes expressed throughout the IDC and changes in chromatin states.

### ApiAP2 transcription factors display distinct associations with chromatin states

Transcription factors (TFs) can both influence and be influenced by chromatin states. In animal genomes, TFs display varied binding preferences depending on both DNA sequence and chromatin environment (Calo & Wysocka, 2013; Srivastava & Mahony, 2020). Some TFs preferentially bind to promoter-associated chromatin states, while others target distal enhancer regions (Grossman et al., 2018; Rabinovich et al., 2008). Certain repressor TFs, such as the repressor element-1 silencing transcription factor (REST), primarily bind in heterochromatic regions (Abrajano et al., 2009). So-called pioneer TFs can actively modify chromatin states by creating regions of chromatin accessibility at their binding sites (Friedman & Kaestner, 2006; Iwafuchi-Doi et al., 2016; Kalyan K. Sinha et al., 2023).

Relatively little is known about the chromatin state associations of TFs in *P. falciparum* (Bonnell et al., 2024). To address this gap, we analyzed the DNA-binding locations (measured via ChIP-seq) of 18 ApiAP2 transcription factors during the IDC. Our analysis revealed that these 18 TFs cluster into three distinct categories (**A-C**) based on the patterns of chromatin states at their DNA binding sites (**Fig. 5**):

**Figure 5:**
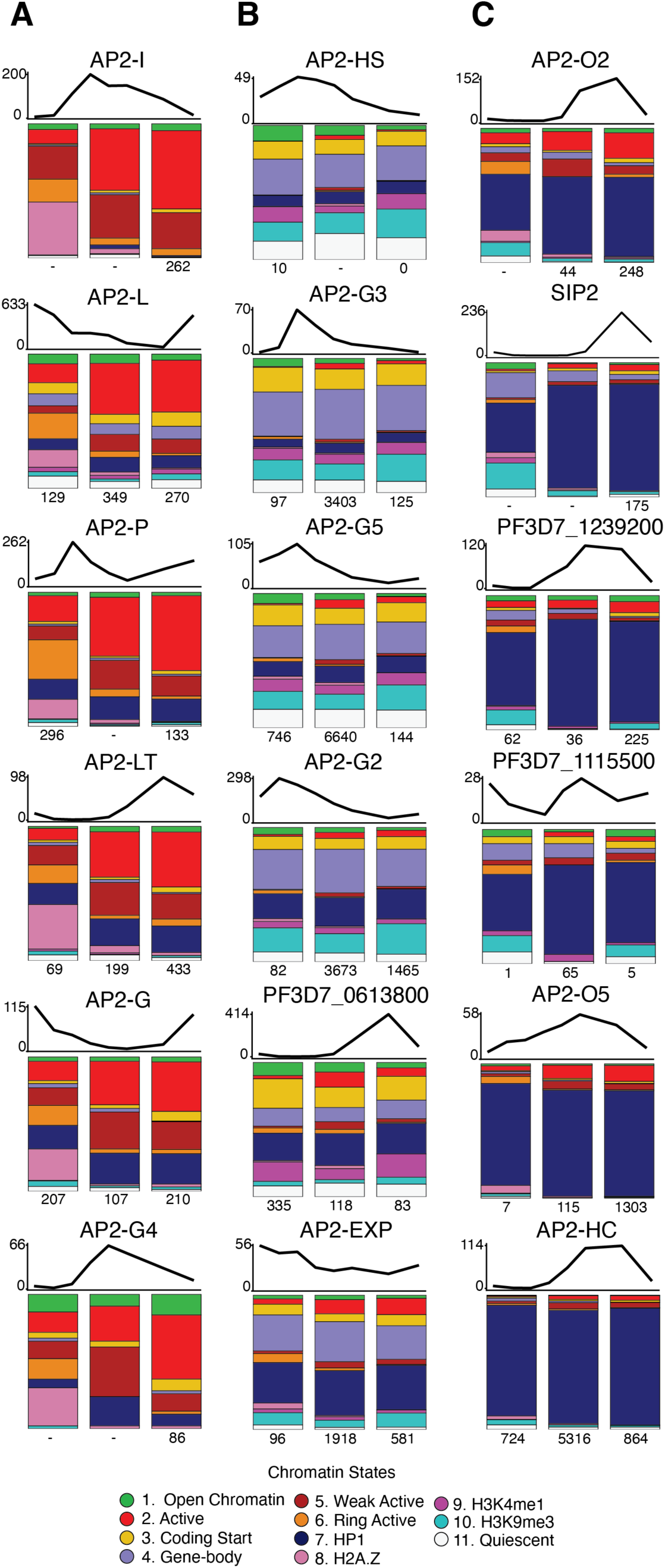
Bar plots show the relative frequencies of chromatin states at transcription factor binding sites across three developmental timepoints (10hpi, 30hpi, and 40hpi). For each transcription factor, all ChIP-seq peak locations were analyzed to determine the chromatin state at the peak summit. Above each set of bar plots, gene expression line graphs indicate the temporal expression pattern of each transcription factor (FPKM values). Numbers below each bar represent the total count of ChIP-seq peaks identified for each transcription factor at the corresponding timepoint. **A)** Transcription factors in the “active” category, which predominantly bind to promoter-associated chromatin states. **B)** Transcription factors in the “poised” category, which associate with a mixture of chromatin states including potentially responsive regulatory regions. **C)** Transcription factors in the “heterochromatin” category, which primarily bind to regions marked by the HP1-associated state 7.

The first category (**Fig. 5A**) comprises six TFs that primarily bind to active, TSS-associated chromatin states. AP2-G and AP2-P exemplify this category; both are transcriptional activators associated with increased expression of downstream genes (Josling et al., 2020; Kafsack et al., 2014; Subudhi et al., 2023). These TFs primarily bind to “active” chromatin states (states 2, 5, 6, and 8) that undergo stage-specific changes in abundance. AP2-I shows the strongest preference for active states 2 and 5 within this group, consistent with its role in regulating genes related to cell division, growth, and invasion (Santos et al., 2017). The chromatin state changes at binding sites for this category of factors largely parallel global shifts in active chromatin states, so it is unclear whether these TFs preferentially bind to specific chromatin environments or directly cause chromatin modifications.

The second category (**Fig. 5B**) includes six TFs associated with a more diverse set of chromatin states, notably including the potentially “poised” state 4 (H3K4me1- and H3K9me3-enriched) and state 10 (H3K9me3-enriched). Several TFs in this category, including AP2-G2 and AP2-G5, have been implicated in regulating the transition from asexual to sexual stage of development. Sexual development (or gametocytogenesis) also takes place in infected red blood cells and requires a 12-day maturation process to produce male and female gametes that will be infectious to mosquitoes during a bloodmeal. Therefore, AP2-G2 and AP2-G5 may act to repress the premature expression of key gametocyte genes (Shang et al., 2021; Singh et al., 2021). The most common chromatin states at binding sites in this category (states 3, 4, 7 and 10) show varied relationships with gene expression: states 3 and 10 do not strongly correlate with RNA abundance, state 4 associates with activation in trophozoites and schizonts, while state 7 correlates with low RNA abundance. Unlike the states preferred by the “active” TF category, the preferred chromatin states in this category maintain relatively stable abundance throughout the time course. Given the mixed association with expression patterns, we characterize this category of TFs as “poised” in the IDC and potentially responsive to developmental or environmental signals.

Finally, we identified six TFs that are predominantly associated with the HP1-associated heterochromatin state 7 that is largely found in subtelomeric regions (**Fig. 5C**). Of particular interest in this group is SIP2, which binds at heterochromatin boundaries in telomeric regions (Flueck et al., 2010). A notable shift in chromatin states at SIP2 binding sites is seen following peak SIP2 expression (**Fig. 5C**), transitioning from predominantly HP1-associated chromatin in trophozoite and schizont stages to a mixed set of states in ring stage (i.e., from 40hpi to 10hpi). This shift is remarkable given the overall stability of the HP1 states across the genome, and may reflect SIP2’s crucial role in activating genes during merozoite formation following the schizont stage (Nishi et al., 2025).

## Discussion

In this study, we comprehensively characterized the chromatin landscape of asexual *P. falciparum* parasites by integrating a collection of diverse epigenomic datasets. Using ChromHMM, we defined eleven distinct chromatin states based on combinatorial patterns of six histone modifications/variants, chromatin accessibility (ATAC-seq), and Heterochromatin Protein 1. Our analysis reveals that chromatin states are highly dynamic throughout the IDC, with approximately two thirds of the genome undergoing at least one chromatin state change between the three timepoints examined. These dynamic changes are closely associated with both lifecycle progression and gene expression patterns, highlighting a complex interplay between chromatin structure and transcriptional regulation.

The largest shift in chromatin states occurs during the transition from the ring stage to the trophozoite stages. This transition involves a global gain of H3K4me3, shifting regions from “Ring Active” state 6 to “Active” state 2 and from “H2A.Z” state 8 to “Weak Active” state 5. These states (i.e., states 2, 5, 6, and 8), and the H3K4me1-associated state 9 are highly dynamic, with frequent transitions to and from other states between timepoints. Meanwhile, the heterochromatin state 7 and the quiescent state 11 are highly stable. This suggest that there are regions of the malaria parasite genome that are stably regulated throughout asexual development versus those that are rapidly remodeled.

Temporal chromatin shifts are closely linked to the parasite’s dynamic gene expression program. While transitions to active, acetylated states (states 2, 3, and 6) at gene start codons correlate with increased transcription and shifts to H3K9me3-marked states (states 4, 7, and 10) generally correlate with expression decline, other transitions revealed more complex relationships. For instance, gaining the “Weak Active” state 5 is paradoxically associated with reduced expression, suggesting a nuanced role in potentially fine-tuning or dampening transcription at highly active loci.

Notably, the HP1-associated state 7 is strongly correlated with transcriptional repression. This state is also the most stable across the IDC, indicating that genes marked by this state are consistently repressed during asexual development. State 7 is enriched at clonally variant genes in sub-telomeric regions, including the *var* (encoding PfEMP1) and *rifin* gene families involved in immune evasion. These genes typically display mutually exclusive expression patterns (i.e., one at a time) within individual parasites (Pickford et al., 2021; Tripathi et al., 2022), explaining their population-averaged association with the repressive HP1 state.

By analyzing ApiAP2 transcription factor binding, we identified three distinct chromatin state associations. ApiAP2 TFs in the “Active” category, including AP2-I and AP2-P predominantly bind to TSS-associated chromatin states (2, 5, 6, and 8). These factors generally regulate invasion and pathogenesis genes (Santos et al., 2017; Subudhi et al., 2023), with the notable exception being AP2-G, which controls gametocyte commitment (Hollin & Le Roch, 2020; Josling et al., 2020; Kafsack et al., 2014; Llorà-Batlle et al., 2020). Their association with TSS-related chromatin states suggests they function primarily as transcriptional activators. While our temporal resolution cannot definitively determine whether these TFs require accessible chromatin or actively remodel chromatin themselves, the expression dynamics of some TFs correlates with the timing of active state transitions at their binding sites.

The “Poised” TF category bind to more diverse chromatin states less strongly correlated with active transcription, including “Gene Body” state 4 and H3K9me3-associated state 10. These TFs may function to attenuate gene expression without fully silencing genes through heterochromatin formation. Several TFs in this category, including AP2-G2 and AP2-G5, are thought to repress gametocyte-related genes (Shang et al., 2021; Singh et al., 2021) potentially allowing for quicker transcriptional activation compared to fully heterochromatic genes.

The “Heterochromatin” category comprises TFs that primarily bind to sub-telomeric regions marked by state 7, suggesting roles in maintaining heterochromatin or regulating clonally variant genes (Flueck et al., 2010; Shang et al., 2022). Our categorization partially aligns with a previous study that defined eight ApiAP2 TFs as heterochromatin-associated factors (AP2-HFs) based on correlation with HP1 binding (Shang et al., 2022). Four of these (AP2-O2, PF3D7_1239200, AP2-O5, and AP2-HC) fall within our heterochromatin-binding category, while three others (AP2-G5, AP2-G2, and AP2-EXP) align with our “Poised” TF category. Interestingly, AP2-P (PfAP2-11A in Shang et al., 2022) falls within the “Active” category in our analysis.

Our findings are complementary to a previous study in which a five-state ChromHMM model was used to investigate a limited set of histone marks (H3K27ac, H3K9ac, and H3K4me3) in egressing *P. falciparum* schizonts, blood cell invading merozoites, and early ring stage parasites (Reers *et al*., 2023). Reers, et al. identified a promoter-associated state that has high signals for all three marks, aligning with state 2 in our study (**Fig. 1**). This state showed a strong association with ring stage expression, even though it is present at relatively few genes in ring stage parasites. Reers, et al. also found that their state 2-equivalent state was highly prevalent in merozoites, a stage for which we did not include data. Another promoter-associated state in the Reers, et al. study, characterized by high H3K9ac and H3K4me3 but low H3K27ac, was consistent with state 5 in our model. This state was prevalent at promoters in schizonts, but less so in merozoites. Additionally, Reers, et al. identified a state with high H3K9ac signals that was enriched in ring stage parasites but not merozoites, which is likely consistent with our state 8. In the IDC stages tested, Reers, et al., did not identify an equivalent to our state 6, which would have high levels of H3K27ac and H3K9ac but low H3K4me3. This difference is possibly due to the higher number of eleven states used by our model, enabled by our inclusion of more chromatin signals. Overall, our study and that of Reers, et al. provide consistent views of chromatin regulatory dynamics at different developmental stages. Our study focuses on the IDC, while they focused on egress and reinvasion, demonstrating the value of a more comprehensive, integrated chromatin state analysis across all stages of the malaria parasite lifecycle.

Although our work offers new insights into the chromatin landscape of *P. falciparum*, several limitations should be noted. The epigenomic datasets originated from different laboratories, introducing potential variability despite our efforts to standardize analysis pipelines. Our analysis is restricted to available histone modifications at three timepoints, potentially missing important modifications or finer temporal dynamics. Future work would benefit from standardized, comprehensive epigenomic profiling across all asexual stages and functional studies to better characterize ApiAP2 TF activities.

Our application of chromatin state analysis to the malaria parasite *Plasmodium falciparum* highlights the dynamic nature of the chromatin landscape during the IDC. These data also define distinct categories of ApiAP2 TFs based on their chromatin binding preferences. Together, these findings indicate that *P. falciparum* TFs not only recognize specific DNA sequences but also show differential preferences for chromatin environments, potentially explaining their target specificity through a combined recognition mechanism. These findings provide a foundation for future studies aimed at understanding gene regulation in this important human pathogen.

## Methods

### Data sources

Nine data types were used to construct chromatin states in this study: ChIP-seq for H3K9ac (Bartfai et al., 2010), H3K4me1 (Tang et al., 2020), H3K4me3 (Bartfai et al., 2010), H2A.Z (Tang et al., 2020), H3K27ac (Tang et al., 2020), H3K18ac (Tang et al., 2020), H3K9me3 (Jiang et al., 2013), HP1 (Carrington et al., 2021; Shang et al., 2022), and ATAC-seq (Toenhake et al., 2018). The HP1 datasets were processed and aligned separately and were then merged internally by ChromHMM.

Most datasets used for chromatin state definition included samples from all three timepoints (10hpi, 30hpi, and 40hpi), with the exception of H3K9me3, for which data were only available at 20hpi and 40hpi. To maintain consistency across all features, the 20hpi H3K9me3 data were used to represent both the 10hpi and 30hpi timepoints in the model. This approach was supported by the high correlation between the available H3K9me3 timepoints (Pearson correlation r=0.992 using 10kb bins; **Supplementary Fig. 3**), suggesting minimal temporal variation in this mark.

The majority of ApiAP2 TF ChIP-seq datasets (AP2-L, AP2-G4, AP2-HS, AP2-G3, AP2-G5, PF3D7_0613800, AP2-O2, PF3D7_1239200, PF3D7_1115500, and AP2-O5) were obtained from Shang, et al. (Shang et al., 2022). Several TFs were profiled by multiple laboratories, and we used merged peak sets in these cases: AP2-G2 (Shang et al., 2022; Singh et al., 2021), AP2-P (Shang et al., 2022; Subudhi et al., 2023), AP2-LT (Bonnell et al., 2024; Shang et al., 2022), AP2-EXP (Russell et al., 2022; Shang et al., 2022), and AP2-HC (Carrington et al., 2021; Shang et al., 2022). Three additional ChIP-seq datasets were also used: AP2-I (Santos et al., 2017), AP2-G (Josling et al., 2020), and SIP2 (GEO ID: GSE296868).

### Data acquisition and preprocessing

A standardized reprocessing pipeline was used for all datasets to ensure consistency. Raw sequencing reads (fastq files) were quality-trimmed using Trimmomatic (v0.39) with parameters SLIDINGWINDOW:4:30 MINLEN:35 (Bolger et al., 2014). Trimmed reads were aligned to the *P. falciparum* 3D7 reference genome using BWA-MEM (v0.7.17) with the -M option and filtered for alignment quality (minimum Phred score of 30) using Samtools (v1.16.1) (Li, 2013; Li et al., 2009). Duplicate reads were removed using Picard’s MarkDuplicates tool (v2.24.1) with parameters REMOVE_DUPLICATES=true and VALIDATION_STRINGENCY=STRICT (The Broad Institute, 2021). After duplicate removal, biological replicates were merged. For ChIP-seq data, peak calling was performed using MACS2 (v2.2.7.1) with parameters -g 2e+7 -q 0.001 --nomodel --shift 0 --extsize 200 (Zhang et al., 2008).

Gene expression data was derived from RNA-seq datasets published by Toenhake et al. (Toenhake et al., 2018), the same source as the ATAC-seq data. RNA-seq reads were trimmed with Trimmomatic (v0.39) using parameters SLIDINGWINDOW:4:30 MINLEN:35, then aligned using STAR (v2.7.10b) with parameters --runMode alignReads --genomeLoad NoSharedMemory --readFilesCommand zcat --outSAMtype BAM Unsorted --alignIntronMa× 10000 --alignIntronMin 30 (Dobin et al., 2013). Gene-level expression counts were generated using HTSeq-count (v2.0.3) with parameters --mode union --format bam --order name --idattr gene_id --minaqual 20 --stranded=no --type exon (Anders et al., 2015), and converted to Fragments Per Kilobase of exon per Million mapped reads (FPKM) values.

The developmental timing in the Toenhake et al. RNA-seq dataset appeared shifted forward compared to another RNA-seq dataset (Chappell et al., 2020). To correct this discrepancy, seven of the eight ATAC-seq and RNA-seq timepoints were shifted back by 5 hours post-invasion. The 40hpi schizont timepoint was not adjusted as it aligned with the latest timepoint in the reference dataset and maintained a consistent schizont stage measurement.

### Chromatin state modeling

ChromHMM (version 1.23) was used to call states, with each developmental timepoint treated as a distinct cell type in the ChromHMM framework (Ernst & Kellis, 2012). For each chromatin feature, BAM files were binarized using ChromHMM’s BinarizeBam function with a bin size of 200 bp and using appropriate control files for each experiment. The LearnModel function was then applied to construct the chromatin state models.

To determine the optimal number of chromatin states, models with 4 to 25 states were generated and evaluated based on several criteria: the percentage of genome covered by the quiescent state; the number of states with similar emission profiles; and the genomic coverage of the least frequent state. As the number of states was increased, the proportion of genome classified as quiescent decreased non-linearly. The “elbow point” in this relationship was identified, beyond which additional states primarily subdivided existing categories rather than identified truly distinct chromatin types (**Supplementary Fig. 4**). By analyzing correlations between emission profiles across models with different numbers of states, it was determined that 11 states optimally captured chromatin diversity while minimizing redundancy.

The relative feature enrichments shown in **Fig. 1B** were calculated using the formula implemented in ChromHMM’s OverlapEnrichment function. This formula computed the ratio of observed overlap to expected overlap (assuming independence between states and genomic features), providing a measure of association strength. The enrichment values were normalized column-wise to highlight which chromatin state showed the strongest association with each genomic feature

### Associations between gene expression and chromatin states

To assess gene expression dynamics and chromatin state dynamics (**Fig. 3**), states were assigned to genes based on the predominant chromatin state near the start codon of the gene. For each gene, the most frequent chromatin state in a 600bp window around the start codon was identified. If more than one state tied for the most overlap, measured in bases, one of the highest tied states was randomly selected.

To associate chromatin states with expression (**Fig. 1E**), each gene was assigned to a state if the gene contained the state or if the state was within 2kb upstream of the gene’s transcription start site (TSS). In cases where two genes were nearby, the closest TSS was used. For overlapping genes, both genes were assigned to the state. The average expression associated with each state was calculated and presented in **Fig. 1E** to represent state-centric expression associations.

### Associations between transcription factor binding sites and chromatin states

For each TF, the union of ChIP-seq peaks across timepoints was taken, and bedtools intersect (v 2.30.0) was used to filter out overlapping peaks to avoid overcounting (Quinlan & Hall, 2010). Pybedtools (v 0.9.0) (Dale et al., 2011) was used to compare TF binding sites to chromatin state locations across all three timepoints. Importantly, chromatin states can still be examined at transcription factor binding sites even when ChIP-seq data is not available for a particular time point, as peak coordinates allow for the analysis of chromatin states at those coordinates across all three time points. All results are displayed as proportions in **Fig. 5**, with each column representing the proportion of the union of TF binding sites covered by each chromatin state at each of the three timepoints. All TF chromatin state associations were determined to be statistically significant by a chi-squared test.

## Acknowledgements

This study was supported by the National Institutes of Health grant R35 GM144135 (to S.M.). A.S.B. gratefully acknowledges funding and training opportunities from the Eukaryotic Gene Regulation training grant T32 GM152354 and from a GlaxoSmithKline Graduate Fellowship. The authors acknowledge Tori Bonnell’s assistance in initiating this study and collecting chromatin data resources from the *P. falciparum* IDC. The authors are also grateful to Till Voss and Richard Bartfai for publicly releasing SIP2 ChIP-seq data prior to publication.

## Availability

BED files containing chromatin state tracks for each timepoint are available at https://github.com/seqcode/pfal-chrom-states-trackhub. A browsable set of chromatin state and associated data tracks are available via the UCSC genome browser: http://genome.ucsc.edu/cgi-bin/hgTracks?db=GCF_000002765.5&hubUrl=https://seqcode.github.io/pfal-chrom-states-trackhub/trackhub/hub.txt

## Supplementary Data

**Supplemental Figure S1.**
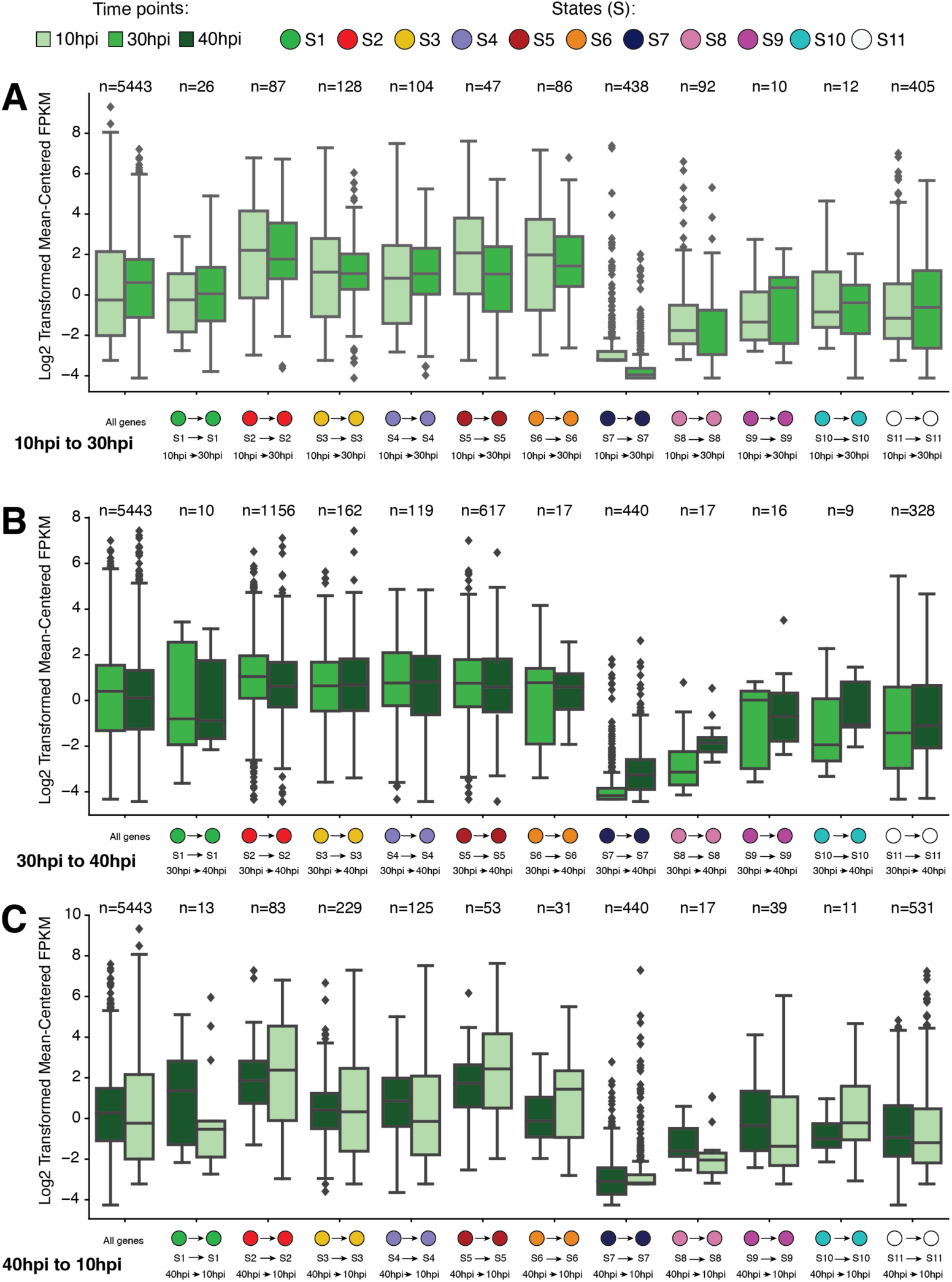
Associations between chromatin state transitions and expression dynamics at genes that do not change states between consecutive pairs of timepoints. Expression boxplots display the distributions of log_2_-transformed mean-centered FPKM normalized RNA-seq values for groups of genes that display indicated self-transitions for each labeled state at the labeled timepoints. **A)** Chromatin state self-transitions and expression values from 10hpi to 30hpi. **B)** Chromatin state self-transitions and expression values from 30hpi to 40hpi. **C)** Chromatin state self-transitions and expression values from 40hpi back to 10hpi.

**Supplemental Figure S2.**
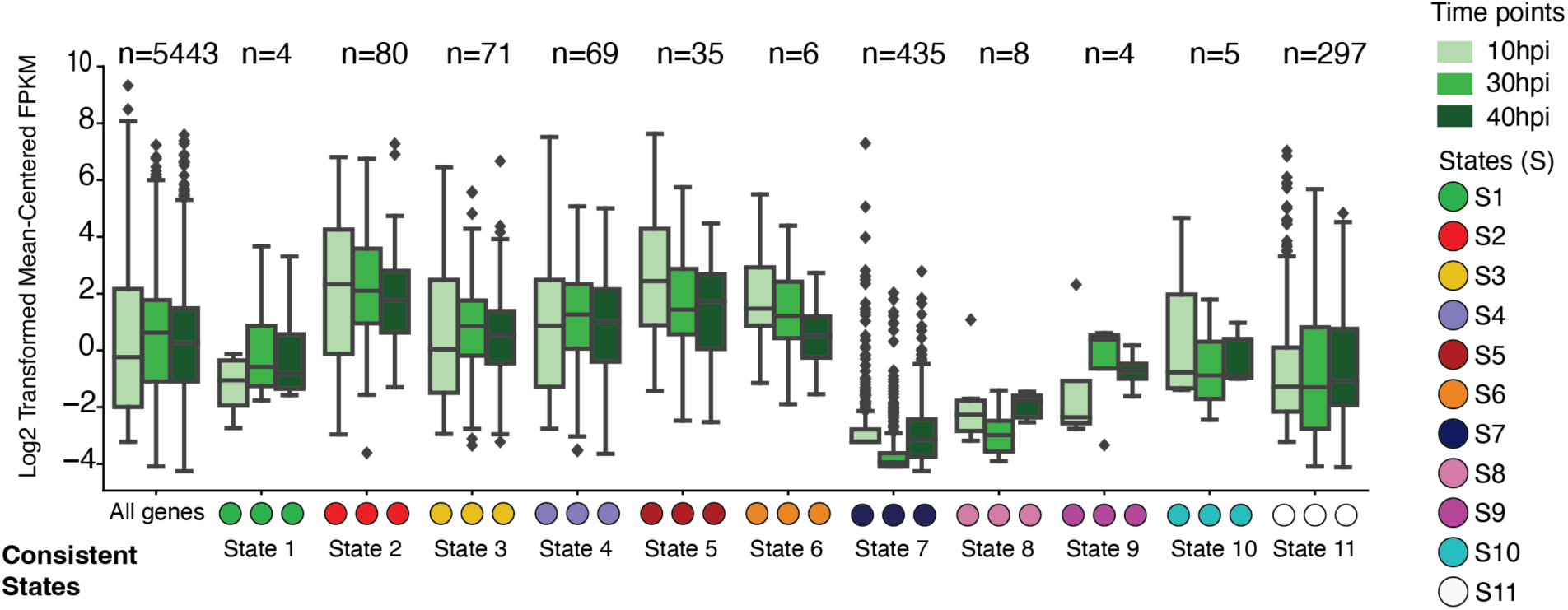
Associations between chromatin state transitions and expression dynamics at genes that never change states across all three timepoints. Expression boxplots display the distributions of log_2_-transformed mean-centered FPKM normalized RNA-seq values for groups of genes that display indicated maintained states for each labeled state at the labeled timepoints.

**Supplemental Figure S3.**
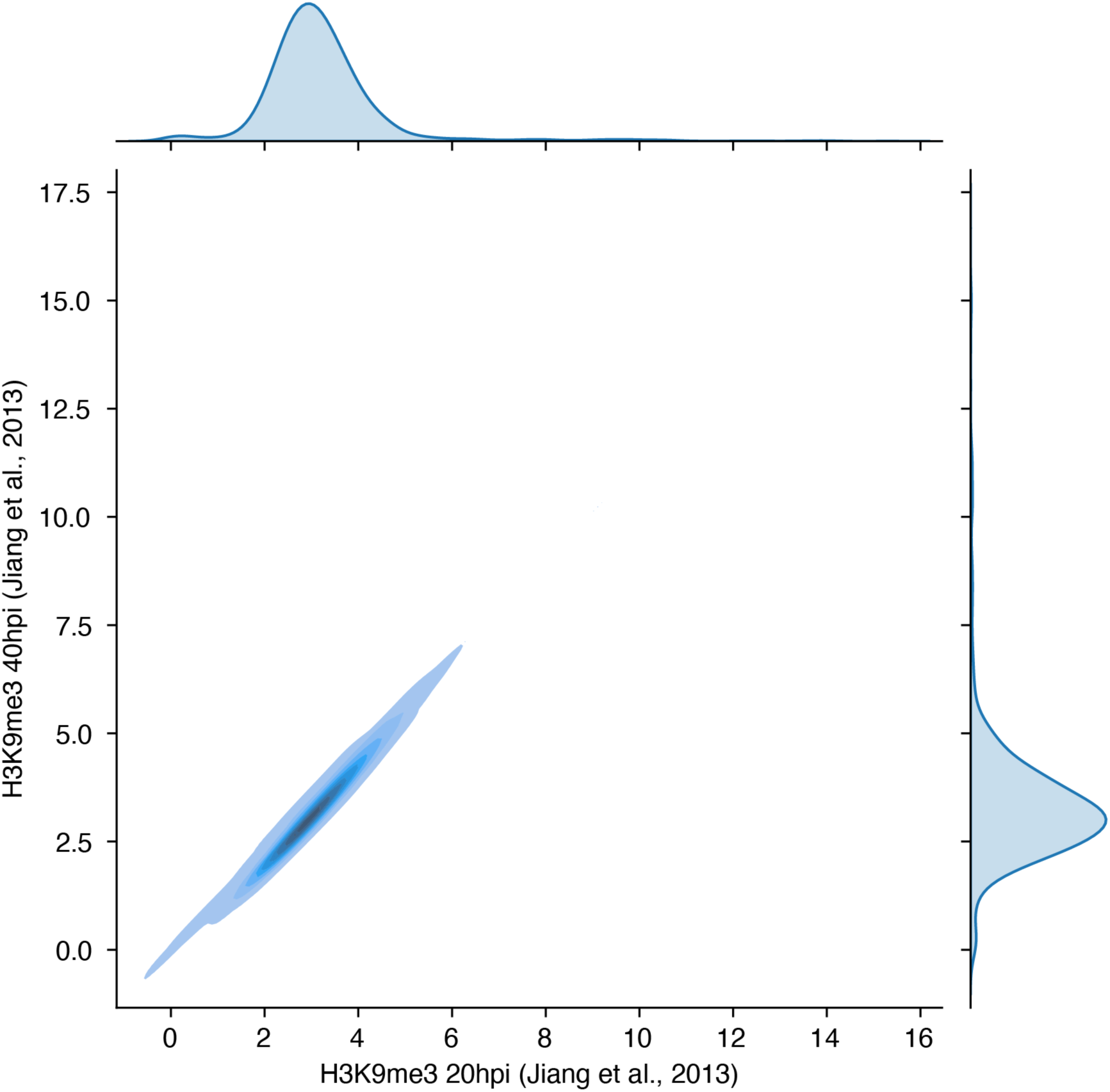
Density plot of binned ChIP enrichment levels between the two H3K9me3 datasets from (Jiang et al., 2013), justifying use of the 20hpi timepoint for both 10hpi and 30hpi timepoints in the chromatin state analysis. Pearson correlation coefficient = 0.992. The density plot and correlation are based on 10kbp bins across the genome using raw read counts.

**Supplemental Figure S4.**
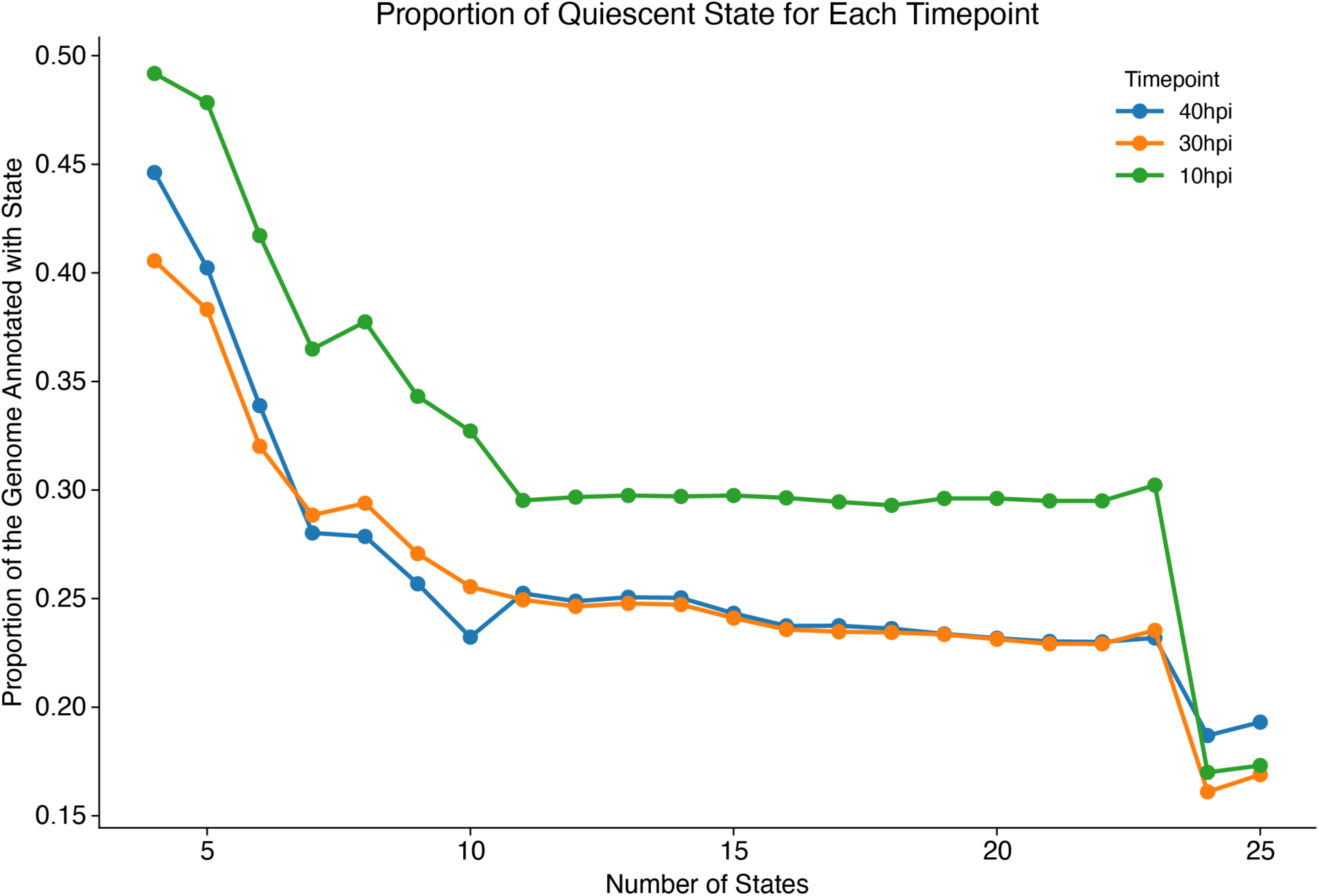
Genome wide proportion of the quiescent state for ChromHMM models varying by the number of chromatin states generated from 4 states to 25 states. All three timepoints are shown for each number of states. The “elbow point” occurs at state 11.

**Supplemental Table 1:** Datasets, references, and accession numbers used in this study.

| **Data** | **Reference** | **GEO ID** |
| --- | --- | --- |
| ATAC-seq and RNA-seq | (Toenhake et al., 2018) | GSE104075 |
| H3K9ac | (Bartfai et al., 2010) | GSE23787 |
| H3K4me1 | (Tang et al., 2020) | SRP252482 |
| H3K4me3 | (Bartfai et al., 2010) | GSE23787 |
| H2A.Z | (Tang et al., 2020) | SRP252482 |
| H3K27ac | (Tang et al., 2020) | SRP252482 |
| H3K18ac | (Tang et al., 2020) | SRP252482 |
| HP1 | (Carrington et al., 2021; Shang et al., 2022) | GSE154840, GSE184659 |
| H3K9me3 | (Jiang et al., 2013) | SRP022761 |
| AP2-I | (Santos et al., 2017) | GSE80293 |
| AP2-L | (Shang et al., 2022) | GSE184659 |
| AP2-P | (Shang et al., 2022; Subudhi et al., 2023) | GSE190497 |
| AP2-LT | (Bonnell et al., 2024; Shang et al., 2022) | GSE184659, GSE212052 |
| AP2-G | (Josling et al., 2020; Shang et al., 2021) | GSE120448, GSE149774 |
| AP2-G4 | (Shang et al., 2022) | GSE184659 |
| AP2-HS | (Shang et al., 2022) | GSE184659 |
| AP2-G3 | (Shang et al., 2022) | GSE184659 |
| AP2-G5 | (Shang et al., 2021, 2022) | GSE184659, GSE149774 |
| AP2-G2 | (Shang et al., 2022; Singh et al., 2021) | GSE184659, GSE157753 |
| PF3D7_0613800 | (Shang et al., 2022) | GSE184659 |
| AP2-EXP | (Russell et al., 2022; Shang et al., 2022) | GSE184659, PRJNA818769 |
| AP2-O2 | (Shang et al., 2022) | GSE184659 |
| SIP2 | Till Voss and Richard Bartfai | GSE296868 |
| PF3D7_1239200 | (Shang et al., 2022) | GSE184659 |
| PF3D7_1115500 | (Shang et al., 2022) | GSE184659 |
| AP2-O5 | (Shang et al., 2022) | GSE184659 |
| AP2-HC | (Carrington et al., 2021; Shang et al., 2022) | GSE184659, GSE154840 |

